# Morphogenesis of the diamond-type stereom microlattice and the origin of saddle-shaped minimal surfaces in the echinoderm skeleton

**DOI:** 10.1101/2025.10.27.684615

**Authors:** Kamil Humański, Giulio Facchini, Jenifer Croce, Dorota Kołbuk, Philippe Dubois, Przemysław Gorzelak

## Abstract

Echinoderm endoskeleton has a unique trabecular microstructure, called stereom, which can exhibit highly ordered geometries. A striking example of these geometries is the recently discovered “diamond-type” stereom, characterized by a diamond triply periodic minimal surface (D-TPMS) and distinguished by exceptional mechanical and structural properties. Despite its promise for engineering applications, the morphogenesis of this microarchitecture remains poorly understood. Here, we applied a multimodal imaging and labeling approach to investigate the developmental processes underlying the formation of the diamond-type stereom in adults of a starfish *Protoreaster nodosus*. We showed that this stereom type develops through two principal marginal growth patterns: trabecular trifurcation, which typically occurs horizontally on flat external plate surfaces, oriented approximately along the crystallographic {111} planes of the D-TPMS; and trabecular bifurcation, which generally occurs along plate edges and between trifurcating zones, aligned with the crystallographic {100} planes of the D-TPMS. Although these growth patterns may proceed at different rates, they are tightly coordinated, producing the coherent D-TPMS microarchitecture. Furthermore, we demonstrated that F-actin cytoskeletal deposition is consistently associated with active biomineralization fronts in both diamond-type and less ordered, galleried stereom. Notably, the formation of lateral bridges between adjacent stereom trabeculae is often preceded by catenoid-like F-actin structures, suggesting a guiding and templating role for the cytoskeleton in building trabecular connectivity and shaping its curvature. Given the recurrence of saddle-like features across stereom types, we hypothesize that minimal-surface geometries may emerge from tension-driven cytoskeletal dynamics acting as physical templates during biomineralization. Our observations underscore the critical involvement of the cytoskeleton in adult echinoderm biomineralization.

**Statement of Significance:** Triple periodic minimal surfaces (TPMS) are geometrically regular, three-dimensionally repeating surfaces that minimize area by maintaining zero mean curvature. Among them, the diamond-type (D-TPMS) is notable for exceptional mechanical stability, uniform porosity, and low surface area-to-volume ratio. Although TPMS-like microarchitectures occur in nature, D-TPMS structures, with large lattice parameters (>10 µm) are exceedingly rare; one natural example is the stereom of certain echinoderms. Yet the mechanism governing the formation of such ordered microstructures remains unresolved. We applied a multi-modal approach to investigate principles of diamond-type stereom morphogenesis. Our results provide key insight into the growth dynamics of this microarchitecture and highlight the essential role of the cytoskeleton, particularly F-actin, in guiding trabecular connectivity and the emergence of minimal-surface geometry, in both periodic and more irregular stereom types.

## 1. Introduction

Stereom is a unique form of skeletal microstructure, regarded as a synapomorphy of all echinoderms [1]. It is a three-dimensional meshwork of trabeculae that branch and intersect, forming a porous and mechanically robust single-crystal magnesian calcite [2], which develops with the involvement of organic molecules during biomineralization [3]. Variations in stereom porosity, regularity, and structural directionality have led to the distinction of several stereom types, including rectilinear (trabeculae arranged in a regular cubic or orthorhombic lattice), galleried (parallel trabecular galleries oriented in a single direction), and labyrinthic (irregular trabecular meshwork), among others [4]. Some of these stereom types have been reported as carrying phylogenetic signals useful for resolving relationships among echinoderm taxa [5]. Furthermore, within the same organism, various stereom types may occur in different skeletal elements or within the same element of a single organism, often reflecting functional specialization of those elements [4-7].

Despite their structural diversity, all stereom types share a common saddle-like profile [4] and display a similar basic developmental pathway [8-12]: their formation begins with the growth of conical protrusions (trabecular tips) that become interconnected through lateral bridging, giving rise to trabecular meshwork. As such, these early structures give rise to a fine meshwork of inner stereom, which may successively thicken and expand at a slow rate through the incremental deposition of biomineral layers. At the cellular level, according to the current model of stereom growth, the elongation of trabeculae is mediated by skeleton-forming cells (referred to as sclerocytes or calcoblasts), which extend thin cytoplasmic processes (skeletal sheaths) that envelop the growing trabecular tips [13, 14]. Between the cytoplasmic processes and the surface of the already deposited biomineral, a confined space occurs in which biomineralization takes place [14].

The conservation of the saddle-shape microstructure in the stereom of all echinoderms suggests that such a special geometry is inherited from a common self-organizing morphogenetic process driving structural complexity. Notably, interactions between soft biological tissues and local curvature have been demonstrated in various organisms and across multiple scales, including the cellular level [15], where they appear to be mediated by the cytoskeleton. Accordingly, in echinoderm embryos and larvae, phalloidin-labeling experiments have recently demonstrated that cytoskeleton plays an important role in biomineralization by guiding morphogenesis. In particular, in echinoid embryos it has been shown that F-actin accumulates around the growing larval spicules [16, 17], particularly at their extending tips [18]. Furthermore, inhibition of F-actin has been shown to induce abnormal, ectopic branching of larval spicules [19]. Recently, F-actin filaments have also been observed connecting pairs of trabeculae in the stereom constituting the sclerites of sea cucumber juveniles (young post-metamorphosis adults), apparently initiating directed growth toward each other, ultimately leading to fusion of opposing trabecular tips [20]. Despite these studies, the role of F-actin in the formation of more complex stereom architectures, particularly within the plates of adult echinoderms, remains largely unexplored. Furthermore, it remains unclear to what extent F-actin is merely a component of a complex skeletogenic machinery, or rather a key determinant of the final curvature profile of the stereom.

In some starfish, sea urchin and sea lily species, the stereom of adults, can exhibit highly organized, regular patterns, such as the recently described "diamond-type" stereom (a specific variant of rectilinear stereom) [21-23]. This structure features a diamond triply periodic minimal surface geometry (D-TPMS) - an exquisitely ordered and periodic surface forming an interconnected diamond-like network, resulting in a highly efficient microarchitecture that is rare in nature [24]. It demonstrates high mechanical strength, including remarkable specific energy absorption capabilities [21]. However, the *in vivo* formation mechanisms of these complex and highly periodic microstructures, especially given their large lattice parameters (>10 µm), as well as the involvement of F-actin during these mechanisms, remain unknown. Indeed, although at the nanoscale, the formation and stability of TPMS with lattice parameters up to ∼100 nm (observed in certain biological systems, including in butterfly wing scales [25]) can be attributed to self-assembly of soft membranes governed by competing molecular interactions among components like lipids or copolymers [26], the mechanism underlying the emergence of such architectures at the microscopic scale remains puzzling. Understanding the biomineralization processes underlying these microstructures would however be of great value, not only to biological sciences, but also to the materials science community.

Here, we explored the growth dynamics and morphogenesis of the recently identified diamond-type stereom in the adult starfish species *Protoreaster nodosus*. By integrating complementary staining techniques with imaging approaches, we characterized the development of this highly ordered microarchitecture. To assess the potential involvement of the F-actin cytoskeleton in stereom formation, we further examine the spatial distribution of F-actin filaments, aiming to elucidate their potential role in shaping the stereom microlattice and regulating biomineralization processes. Altogether, these analyses led us to discover that diamond-type stereom develops through two principal marginal growth patterns, and that the cytoskeleton might play an important role in guiding and templating trabecular connectivity, while also shaping its distinctive curvature profile.

## 2. Materials and methods

### 2.1. Starfish specimens

The study was conducted at the Laboratory of Marine Biology in Bruxelles (Université Libre de Bruxelles, Belgium) on a commercially available (De Jong Marinelife B.V.) and unprotected wild-type (i.e., not genetically altered) starfish species *Protoreaster nodosus*, in which a diamond-type stereom microstructure was recently discovered [21]. Prior to experiments, a total of 25 small starfish, ca. 2.5-6 cm in arm length, underwent a 2-week acclimatization: they were kept in three aerated, closed-circuit, ca. 100 L aquaria containing natural seawater (salinity ∼33.3 psu, temperature ∼27°C, pH ∼7.95) (Fig. S1, Table S2).

### 2.2. Scanning Electron Microscopy imaging

Scanning Electron Microscopy (SEM) imaging was performed on two specimens which were not used in any labeling experiments. A few arm fragments from both specimens were rinsed in ultra-pure water, dried and dehydrated. These samples were then incubated in 5% sodium hypochlorite solution for one hour. A few other arm samples from both individuals were also incubated for 30 minutes in a two-fold diluted solution to preserve the connecting tissue between ossicles and maintain their original spatial arrangement. All of these samples were then rinsed three times in MilliQ water, dehydrated in graded ethanol series and dried. Next, they were coated in platinum and photographed (Fig. 2 and Fig. 3A, B) using a Philips XL-20 scanning electron microscope (SEM) at the Institute of Paleobiology of the Polish Academy of Sciences in Warsaw, Poland (accelerating voltage ca. = 25 kV, working distance ca. = 34 mm). Additionally, a thin section from one specimen illustrated in Fig. 1 was analyzed using a Thermo Fisher Quattro S Environmental SEM (parameters: 20 kV, 0.58 nA, WD = 10mm on carbon coated section) at the Institute of Paleobiology, Polish Academy of Sciences, Warsaw, Poland.

**Fig. 1.**
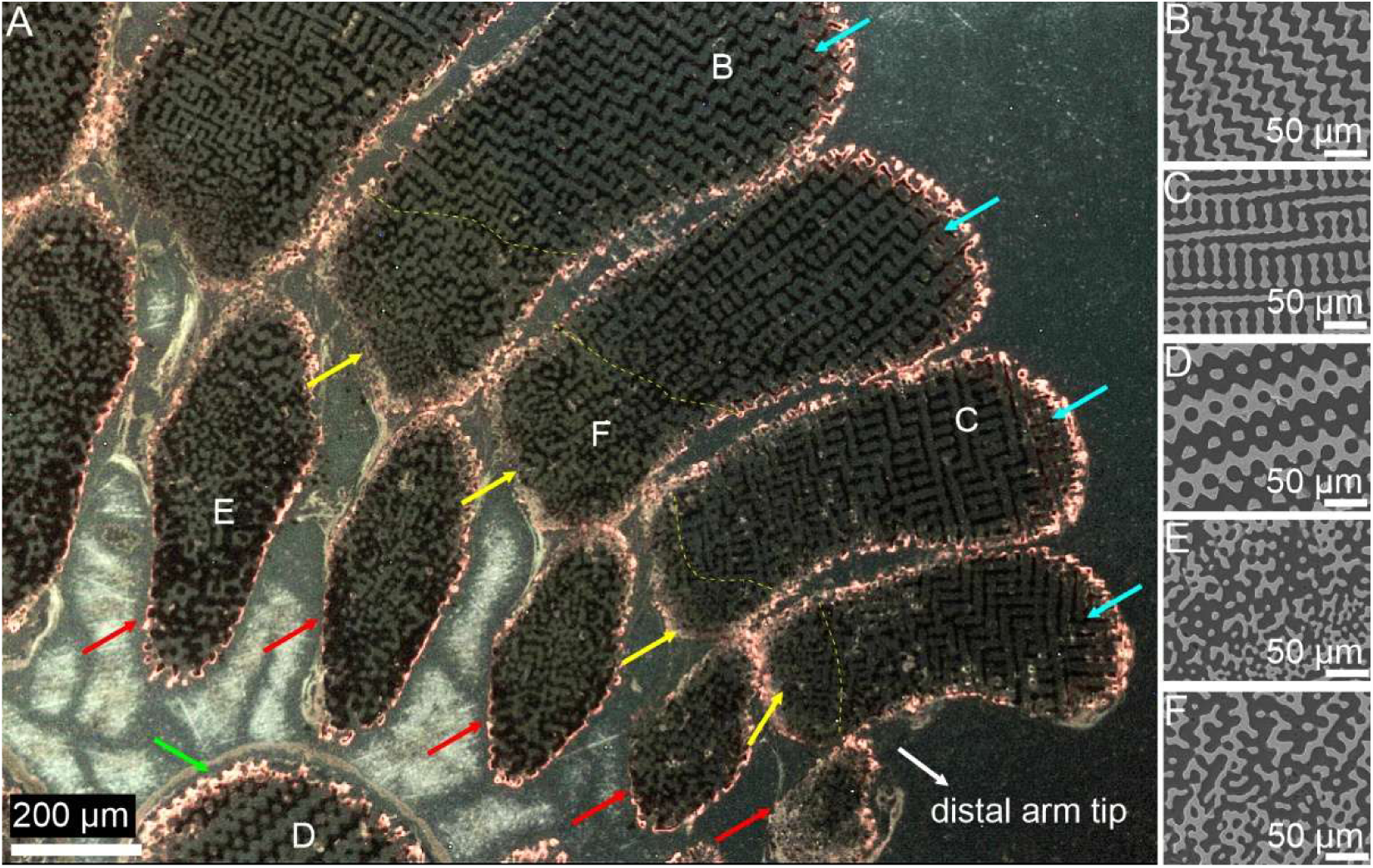
Example of a cathodoluminescence image (A) of a thin section cut along the longitudinal axis of the distal arm portion, with SEM enlargements of selected microregions (B-F). Mn-induced luminescent skeleton appears as a bright orange signal; black and dark grey regions indicate skeleton grown in normal (without Mn^2+^) seawater and resin with soft tissues, respectively. Arrows indicate the following features: green – partially visible carinal plate composed of diamond-type stereom, red – ambulacral plates composed of labyrinthic (unordered) stereom exhibiting regional differences in porosity density, yellow – labyrinthic stereom in the internal portion of the distal adambulacral plates, blue – diamond-type stereom of the adambulacral plates. In the adambulacral plates, the yellow dashed lines indicate the approximate boundaries between the labyrinthic and the diamond-type stereom. B-D show the diamond-type stereom with varying patterns caused by different sectioning planes, while E-F show the labyrinthic stereom [1.5 column image]

**Fig. 2.**
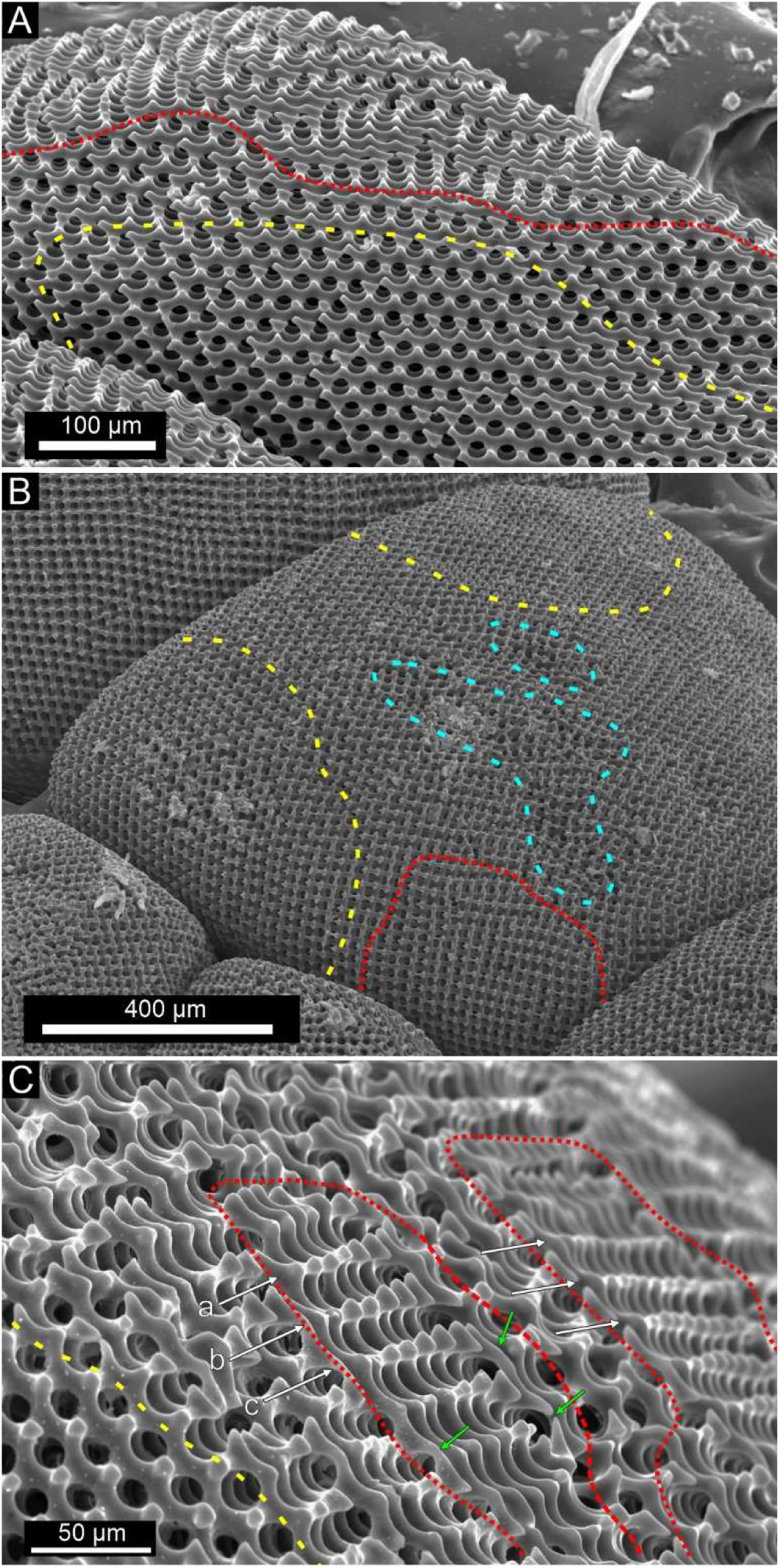
SEM images of superomarginal plates, orientated nearly along the {110} (A) and {100} (B) planes of D-TPMS, and with enlargement of different growth regions on the outer surface of trabeculae (C). Dashed lines highlight distinct growth regions: yellow – approximate extent of trifurcating region; red – approximate extent of bifurcating regions, cyan – a rare occurrence of irregular stereom visible on the surface. The area between red, yellow and cyan boundaries exhibits intermediate growth patterns. White arrows in (C) indicate exemplary bifurcating rows, some of which were denoted (a-c); in each of those a steady progression of growth is observed from a node in the lower left (indicated by arrow) to the node in the upper right. Green arrows mark newly formed trabecular arches (see Discussion). [single column figure]

**Fig. 3.**
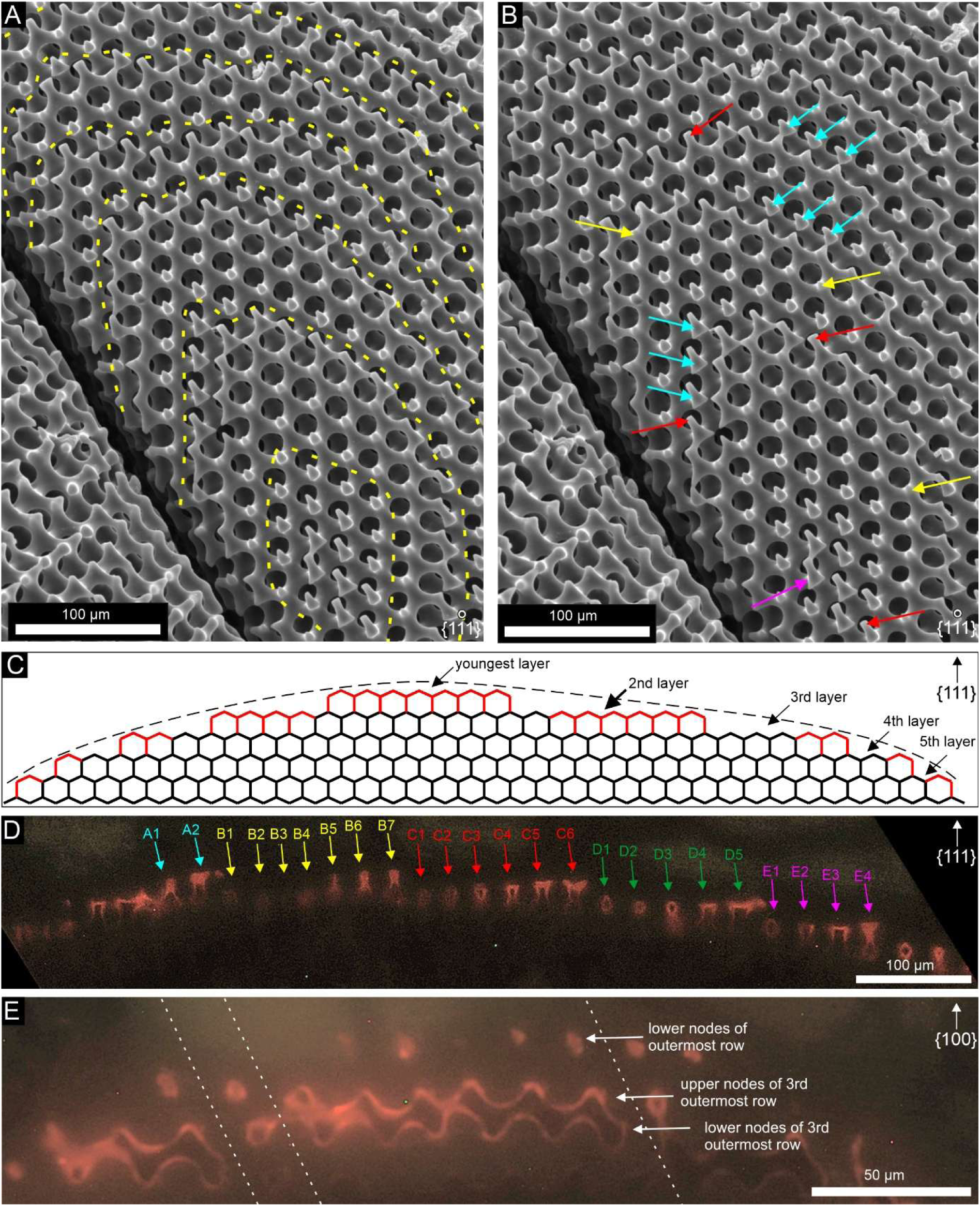
Trifurcating (A-D) and bifurcating (E) regions of the D-TPMS. (A, B) Same region of superomarginal plates viewed along their {111} plane. In (A), yellow dashed lines demarcate the approximate boundaries of successive layers and, in (B), arrows point to the group of nodes forming a new uppermost layer (magenta), to proto-nodes (red), “dormant nodes” (yellow) and crown nodes (cyan) growing at the edges of expanding layers (for definitions see subchapter 3.3). (C) Simplified scheme of trifurcating growth dynamics, based on compilation of data from CL studies. Stereom lattice is depicted as viewed in a thin section perpendicular to the {111} plane of D-TPMS. Growing trabeculae and thickening growth have been omitted. Black branches represent all trabeculae formed prior to the biomineralization of the red-colored trabeculae. The red branches correspond to trabeculae that formed within a short time interval (2-3 days). The thin dashed black line outlines the margin of the growing plate. (D) Surface of an inferomarginal plate, exhibiting an active trifurcating region: thin sections cut roughly perpendicularly to the {111} plane of D-TPMS. Arrows labeled with numbers mark the oldest (e.g., A1, B1) and youngest (e.g., A2, B7) trabeculae located at the respective edges of each highlighted layer. (E) Surface of a different inferomarginal plate, within an active bifurcating region. White lines separate angular unconformities on the thin section. [double column figure]

### 2.3. Manganese labeling

Manganese (Mn)-labeling was conducted following the described procedure [12]. Ten individuals (five controls and five experimental) were placed in independent, aerated 1.5 L aquaria (Fig. S1). Each day, the parameters of the seawater (temperature, salinity and electromagnetic force [emf]) were measured three times. Each day seawater was fully replaced in each aquarium. Specimens were fed every second day with soft tissues of mussels. The experiment spanned up to ten days and involved a total of ten specimens (five controls and five experimental). Of these, six specimens (three experimental and three controls) were subjected to the following sequence: one day in normal seawater, two days of incubation (Mn-enriched seawater for experimental individuals, normal seawater for controls), and a final day of recovery in normal seawater. Two additional individuals (one experimental and one control) underwent an extended post-tagging recovery of three days instead of one. The last two individuals (one experimental and one control) followed the same scheme, with only the experimental individual exposed to a secondary Mn-labeling incubation on days 7 and 8, while the control remained in normal seawater, followed by a 2-day recovery. During labeling, seawater was enriched with manganese using MnCl_2_·4H_2_O (Sigma-Aldrich), dissolved to result in a nominal Mn^2+^ concentration of 3 mg/L.

After the experiments, all animals were fixed, rinsed in seawater and then in ultra-pure water before being air dried for 48 hrs at 50°C. Afterwards, they were dehydrated in graded ethanol series (70%, 90% and 100%, each step lasted 2 hrs), and dried again for 48 hrs at 50°C.

#### 2.3.1. Statistical analyses

Three times per day the parameters of the seawater (temperature, salinity and electromagnetic force [emf]) were measured using a WTW Multi 340i multimeter and a Metrohm pH-meter (826 pH mobile) equipped with a combined glass electrode (Metrohm Primatrode), and calibrated everyday with CertiPUR® buffer solutions pH 4.00 and 7.00 (Merck, Darmstadt, Germany). All emf measurements were subsequently converted to pH values following DelValls and Dickson’s method [27] with TRIS/AMP buffers. Differences between measured water parameters in the different aquaria (Mn-treatment and control) were assessed using repeated measures ANOVA. Significance was set at P < 0.05. Overall, sea water parameters remained comparable between the Mn-exposed and control groups, with significant changes detected only over time, but not attributable to treatment (Table S2).

### 2.4. Cathodoluminescence imaging

Thin sections were prepared from two arms randomly selected from each individual, including both experimental and control specimens. Samples were embedded in epoxy resin, and thin sections were prepared using a series of diamond suspensions and finally carbon-coated. They were analyzed with the aid of Lumic HC5–LM cathodoluminescence (CL) microscope equipped with a hot cathode at the Institute of Paleobiology, Polish Academy of Sciences, Warsaw. Images were recorded using a Kappa video camera. An electron energy of 14 keV was applied and, depending on the sample, beam currents between 0.8 – 0.14 mA were used. Exposure times varied between 1.5 and 10 seconds.

Variably oriented D-TPMS lattices within each plate, with stereom trabeculae cut both parallel to their long axis and at oblique angles, were analyzed (Fig. 1). Given the co-alignment of the calcite *c-axis* with the diamond-type stereom in this starfish species [21], the major crystallographic plane family of analyzed sections were inferred by comparing the observed stereom patterns with those illustrated in [21], cross-checked against model diamond-TPMS sections, and further verified using 2D fast Fourier transform (FFT) analysis and polarized-light microscopy (Fig. S3). Sections oriented approximately along crystallographic {111}, {110} and {211} planes of the D-TPMS were found to be the most informative and were therefore used to assess the growth patterns described below. Thin sections are housed at the Institute of Paleobiology, Polish Academy of Sciences, Warsaw (ZPAL As. 2/Protoreaster01-10).

### 2.5. Calcium, F-actin, nuclei and α-tubulin labeling

A group of six specimens, not previously used in any experiments, were used for labeling of calcium, F-actin, nuclei and α-tubulin. We followed a slightly modified protocol recently published by Vyas al. 2024 [20]. First, for calcium labeling, animals were incubated alive for 24 hours in aerated, independent 1.5 L aquaria, filled with seawater containing calcium-binding fluorescent markers. Two different markers were tested, at a final concentration of 300 mg/L: Calcein Blue (Sigma Aldrich - M1255) and Xylenol orange (Sigma Aldrich - 1086770005), both of which bind to newly deposited calcium. This concentration is comparable to that used in previous study [20], where 250 mg/L was applied. To mitigate the impact of Calcein Blue on seawater pH, sodium bicarbonate solution was added to maintain pH at ca. 8.0. Both markers proved effective in all specimens and provided similar results (see Results section). Following incubation, for F-actin, nuclei and α-tubulin labeling, all animals were rinsed for 30 minutes in aquaria containing fresh seawater. No mortality was observed. For each specimen, two arm tips and two spines (knob-shaped arm tubercles of carinal plates) were collected using a sharp blade. Samples were fixed in 4% paraformaldehyde (PFA) solution for 30 minutes at room temperature (RT). The solution was prepared diluting in MFSW a stock solution of PFA 8% prepared in MilliQ distilled water. Next, all samples were rinsed five times for 5 minutes each at RT using the following proportion of Millipore filtered seawater (MFSW) to Phosphate-Buffered Saline (PBS): 1:0, 3:1, 1:1, 1:3 and 0:1. Samples were then rinsed four times for 5 minutes each at RT in 1x PBST (PBS 1x, 0.05% Tween20), before being pre-blocked for 1 hour at RT in 1x Bovine Serum Albumin (BSA) solution in PBS. For F-actin staining we used Phalloidin (1:100) coupled with four different dyes: iFluor 405, ALEXA Fluor 647, ALEXA Fluor 555plus and ALEXA Fluor 633 (Thermo Fisher Scientific). For nuclei labeling, samples were incubated in 4’,6-diamidino-2-phenylindole (DAPI) (Thermo Fisher Scientific) (1:500). For α tubulin labeling, we used anti-α-tubulin antibodies already coupled with FITC (1:100), as referenced by [20]. For those labeling, upon blocking, samples were incubated overnight at 4°C. All stainings were carried out separately or concomitantly depending on the need. Following labeling, samples were rinsed three times 5 minutes each at RT. Confocal micrographs were obtained using a ZEISS LSM 780 and different magnification water objectives ranging from 20X to 63X, at the Matière et Systèmes Complexes Laboratory UMR 7057 in Paris. Fluorescent labeling was excited using 405, 488, 543 or 633 nm lasers usually acting on two or three different and independent scanning sequences. Z-stack was acquired with an interval of 0.57 µm and was later examined using ImageJ Software [28].

## 3. Results

### 3.1. Microstructural observations by Scanning Electron Microscopy (SEM)

The skeletal organization of *Protoreaster nodosus* comprises several distinct types of plates (Fig. S4). As previously reported [21], we observed that the carinal, inferomarginal, superomarginal, dorsolateral, and adambulacral plates exhibit the same diamond-type (D-TPMS) stereom, whereas the ambulacral plates display labyrinthic stereom (Fig. 1, Fig. S4). All of these plates are further covered by small covering spinelet plates bearing less organized galleried stereom (Fig. S4).

Following [21], the porous lattice of plates with D-TPMS geometry can be described as a network of nodes (points where a variable number of trabeculae converge) and branches (trabeculae connecting two nodes), for which both width and length can be quantified. In the D-TPMS lattice analyzed here, the predominant node type is N4 - a node connected to four branches (Figs. 2-4 and Fig. S4C, D). These nodes are arranged in tetrahedral coordination along the {111} crystallographic direction: the basal vertices of the tetrahedron and the apical vertices extending along {111} (as seen on Fig. 3A, B and Fig. 4).

**Fig. 4.**
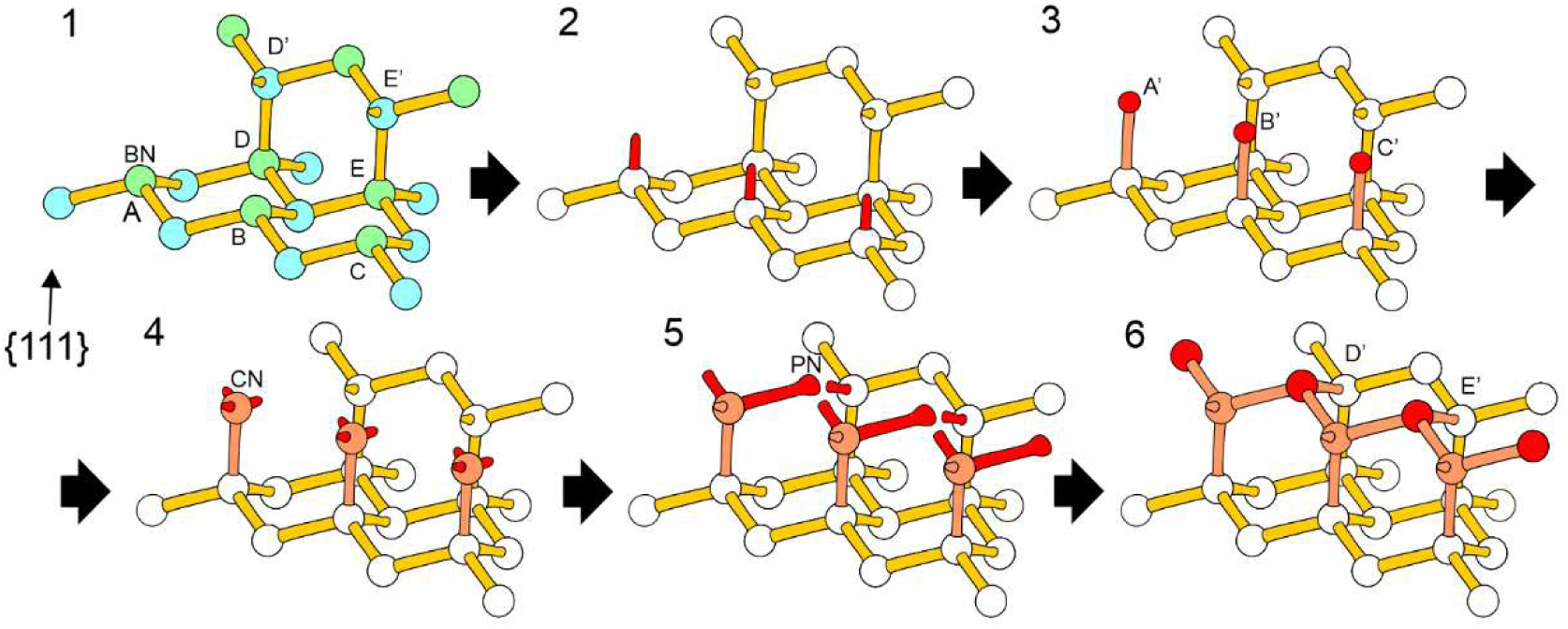
Simplified 3D diagram illustrating successive stages of trifurcating growth localized at the edge of a pre-existing layer, in a plate exhibiting D-TPMS. Nodes are represented with small spheres and branches with cylindrical rods, thus their thickness and length do not represent their actual proportions. Color code: light blue – crown nodes; light green – budding nodes (color distinction shown only in the first panel); yellow – pre-existing branches, that are no longer elongating; white – pre-existing, fully formed nodes; red – nodes and branches undergoing elongation/thickening; orange – nodes and branches that were formed within the depicted time frame, but are no longer elongating. In the figure, we depicted budding nodes from the lower layer (A, B, C, D, E) and crown nodes that grew out of them, belonging to the upper layer (A’, B’, C’, D’, E’), as well as all branches and nodes connected to those ten nodes at the given stage. In the first picture, nodes A, B, C are illustrated at stage 1, nodes D and E are fully developed and nodes D’ and E’ are at stage 6. Other, undefined nodes are fully developed. The symbols BN, CN, and PN were used to denote representative budding node, crown node, and proto-node, respectively. Stage 1. Each budding node (A, B, C) connects only to three crown nodes within the same layer. Stage 2. A budding protrusion emerges upwards along the *c-axis* of calcite from the top of the budding node. Stage 3. A crown node is formed at the tip of the budding protrusion, being noticeably thicker than the budding branch. The original budding node, now connected to four nodes, will no longer develop except for thickening. All further stages refer to the set of newly formed crown nodes (A’, B’, C’). Stage 4. Three protrusions can be distinguished at the top of the crown nodes (around this time, the next set of budding nodes, which are not shown in the diagram, would begin advancing from stage 1 to stage 2). Stage 5. The trifurcating branches elongate and one becomes distinctly longer and thicker at its tip, forming a proto-node (proto-nodes in this diagram are orientated along the layer edge). Stage 6. Branches extending from the crown nodes connect with other adjacent branches, and the developing crown nodes are incorporated into the upper layer. After these connections, nodes D’ and E’ finish their development and only thickening is observed on existing branches. Similarly, nodes A’, B’ and C’ finish their development only after the next set of crown nodes reaches stage 6. Stages 1-3 describe three lower-layer nodes (A, B, C), while stages 3-6 correspond to crown nodes (A’, B’, C’). Proto-nodes, which have connected to two adjacent branches emerging from the nearby crown nodes, form new budding nodes in the upper layer. The minimal estimated (based on CL images) time for progression from a budding node at the beginning of stage 2 to a crown node at the end of stage 6 is approximately 12 hours. [two column figure]

SEM imaging allowed us to study the spatial distribution of skeletal structures and stereom microarchitectures preserved at different developmental stages (Figs. 2-5). Through high-resolution SEM imaging of plate surfaces, we observed repetitive stereom structures associated with distinct patterns of stereom growth, consistently appearing on all D-TPMS-bearing plates. However, elucidating the formation mechanisms of these structures required integrating data from both SEM and cathodoluminescence studies (see subchapter 3.3).

**Fig. 5.**
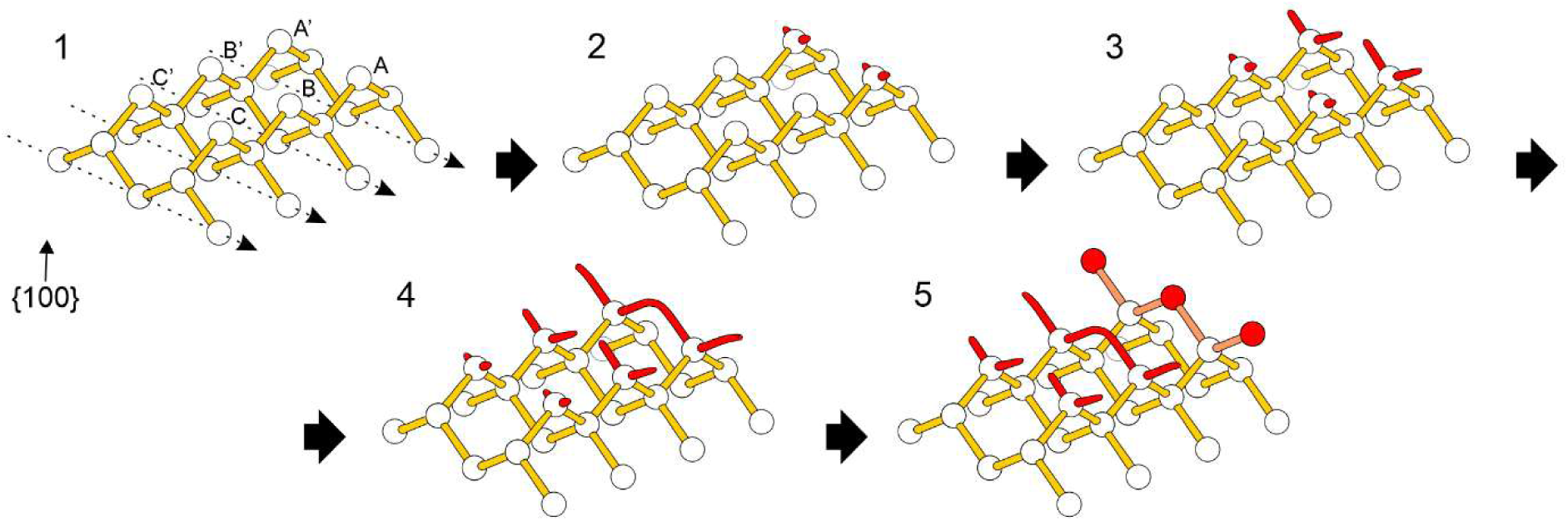
Simplified 3D diagram showing sequential stages of bifurcating growth across three node generations in a plate exhibiting D-TPMS. Node and branch thicknesses and lengths are not representative, as explained in Fig. 4; colors follow those of Fig. 4. Stage 1. The upper nodes in two parallel rows (A-C, A’-C’) are connected to two bottom nodes in the same row. Here, six rows are illustrated: four lower (marked by arrows), and two higher, perpendicular to them (containing nodes A–C and A’–C’). Stage 2. Two side branches emerge from two oldest upper nodes (A, A’), forming a straight top ridge perpendicular to the row. Stage 3. Branches extend and, while growing, incline slightly toward adjacent nodes, while at the same time the subsequent upper nodes start stage 1. Stage 4. At least one branch connects with that of a neighboring node, forming an arch-shaped skeletal bridge. Stage 5. A new node develops from the top of the arch. The new node enters stage 1, starting the next cycle. Nodes B, B’, C, C’ and subsequent pairs (not shown on the diagram) continue to grow until, one after another, they reach stage 5. In this way, growth waves propagate along each row, generating surfaces of parallel node series. Growth typically proceeds at a steady pace during stages 1–2, then accelerates within stage 2. Longer bifurcating series show most nodes at stages 1 or 2 (Fig. 2C - therein rows a-c contain 2-3 such nodes each, then row c includes a stage 4 node; rows a and b reach stage 5) [Two-column figure]

On every plate exhibiting D-TPMS stereom lattice, three main patterns of diamond-type stereom growth were observed (see Fig. 2): (1) here termed trifurcation, which is a stereom growth resulting in the formation of successive layers roughly diamond to elliptical in shape and occurring on flat external surfaces of plates oriented approximately along the {111} planes of the D-TPMS (Fig. 2 and Fig. 3A). (2) here termed bifurcation, which is a growth pattern that appears to be supplementary to trifurcation, typically occurring around the edges of external surfaces of plates or between trifurcating regions on surfaces of varying shape, roughly aligned with the {100} planes of the D-TPMS lattice (Fig. 2 and Fig. 3E); (3) an intermediate growth pattern, which is present at the borders between tri- and bifurcating regions, and exhibits a combination of characteristics from both growth types (Fig. 2A, B). It is worth noting that if the D-TPMS microarchitecture ultimately results from marginal (surface-driven) growth, the range of viable geometric configurations becomes inherently limited, which is consistent with the emergence of bifurcating and trifurcating growth patterns along defined orientations.

Layers in trifurcating regions consist of nodes extending laterally and apically. Each node in lateral vertices (hereafter referred to as crown node) connects to three nodes in apical vertices (hereafter referred to as budding nodes) of the same layer and one budding node in the layer below, while each budding node connects to three crown nodes in the same layer and one crown node in the layer above (Fig. 4). During growth, crown nodes form on top of the branch extending from a budding node, and branches extending from the crown node connect with each other, forming new budding nodes while maintaining the tetrahedral connectivity. This process will be explained in detail in chapter 3.3.

All aforementioned plate types exhibiting D-TPMS were found to further contain small volumes of irregular, unordered labyrinthic stereom (Fig. S5). These regions are typically located within the interior and exhibit ellipsoidal shapes. They are clearly distinct in both size and trabecular thickness from the D-TPMS stereom (which usually displays some minor dislocations [21]). In adambulacral plates, the labyrinthic stereom is located in the internal portion of the plate. The morphogenesis of the labyrinthic stereom has not been explored here.

### 3.2. Cathodoluminescence analysis of stereom growth patterns

Manganese (Mn) labeling followed by cathodoluminescence (CL) imaging has previously been shown to be an efficient method for tracking stereom growth dynamics at the microstructural scale in echinoderms [12]. In adult starfish, Mn-stained stereom was consistently well-differentiated under CL, in all thin sections taken from the arm fragments of specimens in the main experimental group (Fig. 1), with staining being detected in every marked specimen. In contrast, the stereom of the control group showed no Mn-induced luminescence (Fig. S6).

The skeleton labeled with Mn exhibits distinct, bright orange luminescent increments at the plate periphery, indicating that skeletal growth and Mn incorporation occurred during the two-day incubation treatments. In contrast skeletal growth during the one-day recovery period, in the main experimental group, was rarely observed in the examined plates, indicating a delay in biomineralization upon Mn^2+^ exposure, likely caused by the equilibration of manganese pools between ambient seawater and internal tissues of starfish. Note that a single individual incubated for an extended (3-day) recovery period exhibited somewhat more pronounced and clearly detectable non-luminescent skeletal growth following the tagging event (Fig. S7). In contrast, another specimen subjected to two separate Mn-labeling events yielded inconclusive results regarding non-luminescent skeletal growth, as no clearly distinguishable two growth increments were consistently observable.

Precise quantification of growth rates was hindered by orientation-dependent variability of the sectioned surfaces, which introduced measurement bias. Nevertheless, the rate of progression of both tri- and bifurcating growth fronts appeared to vary with plate type and anatomical position within the arm, ranging from several to tens of micrometers per day, and were highly dependent on factors such as the individual specimen, plate size (or inferred age), and specific location on the plate. Only on some surfaces a continuous, evenly thick stereom (approximately twenty micrometers thick) was formed during labeling (Fig.1) while on other plates the Mn-induced luminescent growth fronts contained gaps (Fig. 3D). Thickening of previously formed stereom trabeculae (including both longitudinal trabeculae and lateral bridges) was also well visible, and could extend up to several tens of micrometers below the areas where trabecular tips were forming. In general, the peripheral growth was most pronounced in the distal plates.

Despite these uncertainties, we were able to identify a general growth pattern and formulate hypotheses concerning its dynamics. For instance, if we define the growth rate as the change in the trifurcating layer radius over the measured time interval, one observes that the average growth rate generally decreases towards the lower layers (Fig. 3C, D), relative to the ring thickness, in order to maintain the surface curvature. Considering the minimal size differences between layers, the minimal observed growth rate was ca. ∼15 µm/2 days (corresponding to the incorporation of a single new crown node within that time frame). The maximal observed growth rate of a pre-existing (second youngest) layer in one direction was approximately 40 µm/2 days, equivalent to 3 newly incorporated crown nodes (Table. S8). Conversely, defining the growth rate globally, as the number of new nodes (red in Fig. 3C, D) along a line perpendicular to the dashed envelope, showed that it remains approximately constant (one node per period) within the resolution of our measurements, explaining the preservation of the overall dome shape.

### 3.3. Stereom growth patterns

As mentioned before, the information obtained through SEM was complementary to CL imaging, with SEM providing high-resolution details of stereom architecture, while CL allowed the determination of the sequence of stereom formation (see also Fig. S9). The following subsections present these observations in detail, integrating data from both techniques - collected from five individuals subjected to Mn labeling and two individuals examined using SEM.

#### 3.3.1. Trifurcating stereom growth

Trifurcating growth was only observed on external surfaces of plates oriented along the D-TPMS planes belonging to the crystallographic {111} family. Most prominently, it occurred when the surface of the plate was oriented at an angle between 0° and 30° to the {111} plane, around which it became less distinguishable and was progressively replaced by bifurcating growth mode, which became most pronounced at the plate surface inclined at an angle 30° to 45° relative to the {111} plane. When viewed from the top (Fig 3A), along the {111} plane of the D-TPMS, the stereom formed by this mode of growth appeared as a bundle of layers stacked upon one another (Fig. S9B). The outermost layer in this view, corresponding to the trifurcating region, represented the youngest stage of growth.

As pointed out above, each fully developed crown node is connected to three budding nodes within the same layer and one budding node from the layer below. Conversely, each fully developed budding node is connected to three crown nodes within the same layer and one crown node from the layer above (Fig. 4 Stage 1). When such a layer reached a certain width, few budding nodes located near its center (presumably the oldest from the previous generation) - simultaneously formed small protrusions (buds) that elongated vertically along the crystallographic {111} direction, which corresponds to the *c-axis* of the constituent calcite at the atomic scale (Fig. S3), giving rise to vertical branches directed upwards (Fig. 3B, Fig. S9 and Fig. 4 Stage 2). At the tip of each branch, a new crown node formed (Fig. 4 Stage 3). These new crown nodes then trifurcated (Fig. 4 Stage 4), creating three small lateral protrusions. From each new crown node, one of these protrusions was growing visibly faster (Fig. 4 Stage 5) and eventually formed at their tip a node-like structure, hereafter referred to as a proto-node. New proto-nodes in a given trifurcating region consistently emerged along the same direction (Fig. 3B and Fig. S9). Depending on their location, proto-nodes can thus grow toward (e.g. Fig. 3: lowermost red arrow and Fig. S9, blue nodes in the middle), outward (Fig. 3: uppermost red arrow) or along the layer margin (Fig. 3: two other red arrows), with the latter case being most common on all plates due to the D-TPMS stereom geometry. Subsequently, if another branch or branches from a nearby crown nodes are available for coupling, they connected to the proto-node that ultimately became a new budding node (Fig. 4 Stage 6).

After forming its core, each layer expanded peripherally by incorporating additional crown nodes that budded from the underlying layer in a circular pattern around the growing layer (Fig. 3B). Thus, zones of active trifurcating growth formed concentric rings around the periphery of each layer, spreading outward horizontally like ripples, as the layer increased in size. The horizontal growth of each layer remained approximately isotropic until it approached the plate edge where a non-trifurcating region begins (see Fig. 2A). Budding nodes located between active growth zones remained “dormant”, exhibiting only thickening growth of trabeculae until they reached their maximum width. In general, each successive pair of layers exhibited a decreasing size difference, leading to progressively thinner “rings” of exposed nodes in each subsequent layer (Fig. 3A, C). The minimal ring width (∼30 µm) was observed when it consisted of a single line of exposed budding nodes. This is consistent with marginal growth, occurring mainly at the edges of the 3D structure of the ossicle, i.e., normally to the dashed envelope sketched in Fig. 3C.

The active, thickening, and dormant zones can be observed on CL images (Fig. 3D, E). In these images, the manganese signal was most prominent on the edges of each layer, where nodes and branches were composed entirely of manganese-induced luminescent calcite (active zones) (e.g. Fig 3D, nodes B6-B7 and C5-C6). Few preceding nodes in each layer might also exhibit distinct luminescence, but only on their outer surfaces, which indicates that these nodes underwent only thickening growth during the staining period (thickening zones) (e.g. Fig. 3D, nodes C3-C4). The remaining exposed budding nodes of each layer showed dull or no luminescence, suggesting that no growth occurred during that time (dormant zones) (e.g. Fig. 3D, nodes B1-B4 and C1-C2). The number of those nodes in a layer and the time they spent in “dormant” phase varied depending on the shape of the plate and the size of the layer, however, this duration, based on CL images, can be approximated to range from a day to a week.

#### 3.3.2. Bifurcating stereom growth

As previously mentioned, bifurcating growth is considered supplementary growth, occurring on surfaces orientated at ca. 30-45° angle to any of the {111} planes, where uniform trifurcation is not supported. In contrast to trifurcating regions, which possess a fixed central point, concentric growth zones surrounding it and distinct layers, bifurcating regions exhibited a seemingly irregular growth that only locally exhibited repeating structures (Fig. 2C). As such, bifurcating growth regions could have only been seen as a patchwork of linear rows of different length, which grew by extending one or both of their ends.

The bifurcating growth pattern was most clearly observed in two adjacent, parallel rows as sketched in Fig. 5. Growth initiated from the oldest budding nodes in each row (Fig. 5 Stage 1), giving rise to small protrusions (Fig. 5 Stage 2), which subsequently extended into branches (Fig. 5 Stage 3). As the two branches elongated, they further gradually inclined toward each other (Fig. 5 Stage 3) and as a result connected to each other (Fig. 5 Stage 4). Upon connecting, an initial arch-like structure lacking a distinct central node was formed (Fig. 2C (green arrows), Fig. S9D and Fig. 5 Stage 4). Ultimately, a new node formed upward from the apex of this arch (Fig. 5 Stage 5).

### 3.4. Confocal observations - overall characterization

To gain insights into the potential involvement of the cytoskeleton in stereom growth, regardless of stereom type, and in particular at the level of individual trabeculae and their fusion processes, we next performed a series of staining targeting F-actin filaments and microtubules. Along these staining, we further labeled nuclei and newly biomineral deposition of calcium, to concomitantly visualize trabecular growth. We found, as previously reported [10, 20], the accumulation of calcium located at the growing tips of trabeculae (Figs. 6, 7).

**Fig. 6.**
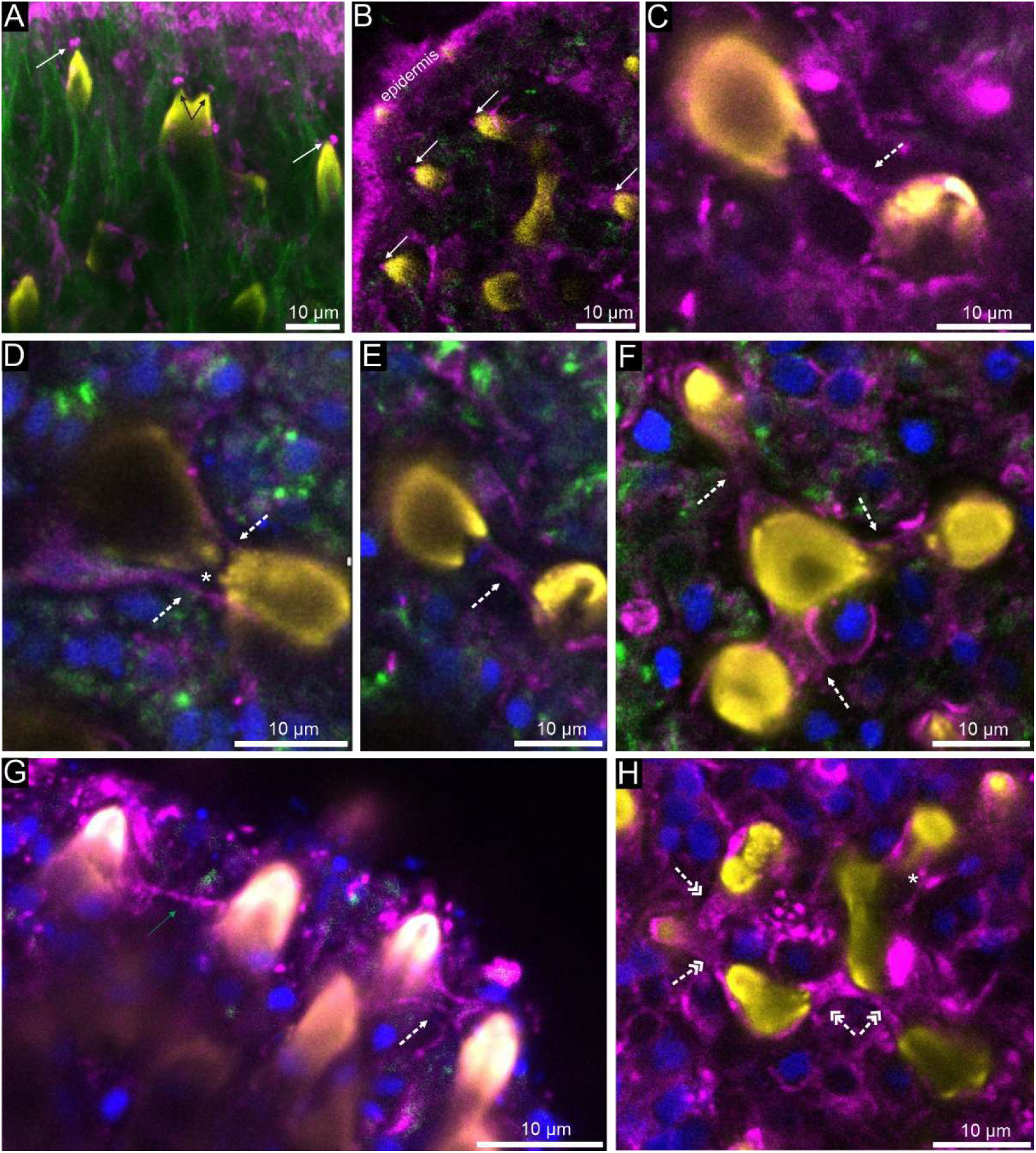
Confocal images of galleried stereom in spinelet plates of an adult starfish showing F-actin densification in growing trabecular tips and between trabeculae. (A, B) Zoom-ins of F-actin caps located at the tip of growing trabeculae. (C-H) Zoom-ins of F-actin catenoid-shaped filaments connecting adjacent growing trabeculae or wrapping adjacent trabecular bridges. Yellow – freshly biomineralized skeleton, blue – nuclei, magenta – F-actin, green – α-tubulin, white plain arrows – F-actin caps, black arrows – trabecular tips, dotted white arrows – F-actin catenoid bridges, * – sheath-like coating of trabeculae, dotted white double arrows – complex F-actin catenoid bridges, green arrow – thin F-actin filament. [1.5 column figure]

Furthermore, we observed that cell nuclei were commonly localized near the base of the growing trabeculae (Fig. 6), rather than at the growing tips, and showed a loosely dispersed pattern therein with no evident spatial regularity. Microtubules, in turn, appeared arranged in bundles, predominantly oriented longitudinally along the trabeculae (Fig. 6A). However, although the α-tubulin signal was typically strong at the onset of image acquisition, it exhibited thereafter rapid deterioration upon increased laser power and depth imaging, impairing further analyses. By contrast, cytoskeletal labeling carried out for F-actin, revealed a similar spatial distribution and three-dimensional organization of F-actin filaments on the growing surfaces of both the diamond-type stereom-bearing plates (carinal, inferomarginal, superomarginal, dorsolateral and adambulacral) and the less-ordered galleried stereom-bearing covering spinelet plates (Figs. 6, 7). A consistent pattern emerged across all observed plates and stereom types in all six marked specimens: F-actin densification was localized at the tips of actively growing trabeculae (identified by strong calcium-related fluorescence signal) regardless of plate morphology or underlying stereom microarchitecture.

In the galleried stereom of spinelet plates, the intensified F-actin signal appeared as minute caps positioned a few micrometers above the tips of growing trabeculae, or as catenoid-shaped structures forming between adjacent elongating branches (Fig. 6). In diamond-type stereom, F-actin signals were also detected at the tips and in between trabeculae, where they form several recurring structural morphotypes (ranging from thin, disordered bundles to catenoid-like tubular structures, and sheath-like coatings), which seem to be related to specific developmental stages of the associated trabecular tips (Fig. 7).

**Fig. 7.**
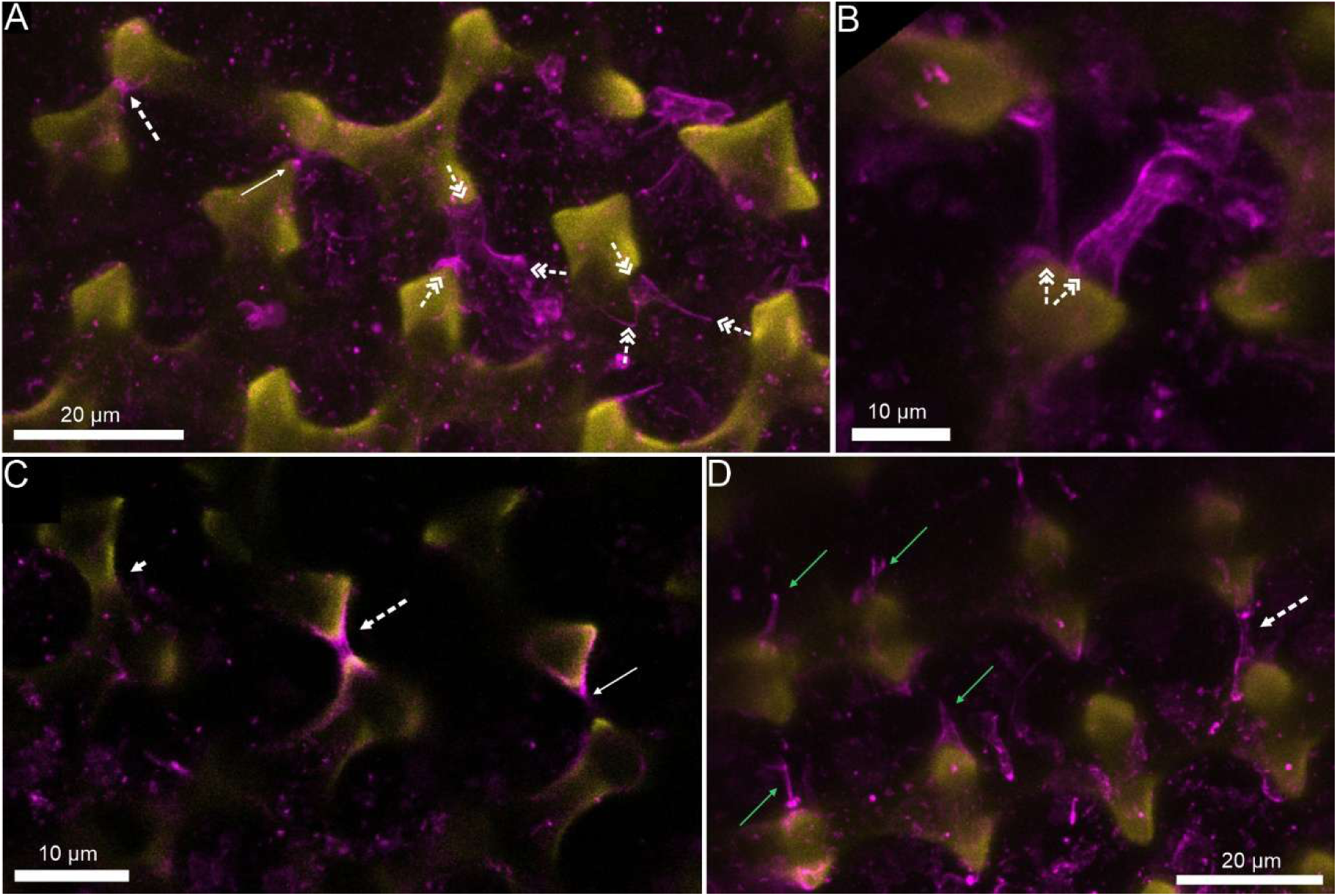
Confocal images of F-actin densification in growing trabecular tips in active growth zones of diamond-type stereom in adambulacral plates of an adult starfish. (A, B) Trifurcating zones. (C, D) Bifurcating zones. Arrows highlight a variety of F-actin structures possibly at different stages of development. Yellow – freshly biomineralized skeleton, magenta – F-actin, white plain arrows – F-actin caps, dotted white arrows – F-actin catenoid bridges, white plain short arrow – fully developed trabecular calcite bridge without F-actin coating, dotted white double arrows – complex F-actin catenoid bridges, green arrow – thin F-actin filament. [2 columns figure]

## 4. Discussion

As noted recently [21], the formation mechanisms of diamond-type stereom have not been systematically explored, and understanding these processes would be of great value not only to the materials science community, but also to a broader audience across multiple scientific domains. Here, we provide the first detailed characterization of these morphogenetic processes, demonstrating that ossicle formation bearing diamond-type stereom can be described in terms of two fundamental growth modes: trifurcation and bifurcation. While progressing on different distinct planes, these growth modes are synchronized, ultimately generating a coherent D-TPMS lattice. Trifurcating growth can be divided into six identifiable stages, as visualized in Fig. 4; it should be noted that the sequence is qualitative and does not imply a constant growth rate or a uniformly distributed timeline. Similarly, bifurcating growth can be described in terms of five observable stages, depicted in Fig. 5, with the same caveat regarding the qualitative nature of the sequence.

Building upon the delineated growth patterns of ossicles, we further explored the cytoskeletal dynamics that is linked to trabecular formation. We found that intensified F-actin signals at the tips of growing trabeculae were consistently observed at active growth fronts in both diamond-type and less-ordered galleried stereom of *Protoreaster nodosus* (Figs. 6, 7). These findings corroborate some earlier reports documenting F-actin accumulation at biomineralization sites in echinoid embryos and holothuroid juveniles [17-20], while representing the first visualization of F-actin distribution in asteroid skeletal elements. The possibility that F-actin participates in initiating or guiding trabecular growth and connections within echinoderm stereom has only recently been proposed [20] and still requires more thorough documentation. Thus, the observations presented here not only support the hypothesis of a key role for the cytoskeleton in echinoderm biomineralization [18-20], but also suggest actin-mediated structural templating/guiding in the form of caps and catenoid-like bridges during stereom morphogenesis.

In general, F-actin structures associated with elongating trabeculae can be classified into two categories: (i) caps, which are situated immediately above or closely associated with growing trabecular tips, and (ii) bridges, which connect two opposing growing trabeculae. Both categories exhibit some variability in shape and size, likely reflecting differences in developmental stage and growth dynamics.

F-actin accumulation was consistently observed across all skeletal plates, irrespective of whether they contained galleried stereom (e.g., spinelet covering plates) or diamond-type stereom, the morphogenesis of which was explored in detail herein. In both stereom types, F-actin caps appear as loosely hemispherical or open conical concentrations, typically thinner than the underlying trabecular protrusions and positioned up to about a few micrometers above the trabecular tips (Figs. 6, 7). Notably, thread-like projections composed of membrane and F-actin forming between the tips of growing trabeculae have recently been documented in juvenile holothurians [20]. Those projections were notably occurring only between approaching trabeculae, suggested to enable their curved growth and eventual fusion [20]. In our study, we demonstrate that within these projections, which were also distinguished only between approaching trabeculae, F-actin forms however structurally distinct cytoskeleton bridges, which likely guide and template the ongoing biomineralization of the stereom lattice. We observed F-actin in various forms, thus potentially in different organizational stages, ranging from thin, disordered bundles (Fig. 6G) to catenoid-like tubular structures (Fig. 6C, H and Fig. 7C), and, hypothetically, even to a sheath-like coating (Fig. 6D, E) filled with newly deposited mineral. Furthermore, we observed complex catenoid-like bridges between multiple neighboring trabeculae (e.g., Fig. 6H). In trifurcating regions of D-TPMS, specifically between proto-nodes extending from the crown node (outward to the layer margin) and two nearby budding nodes, these bridges exhibit a “pants-like” morphology (Fig. 7A, B).

Together, these observations provide new insights into the dynamic processes underlying the formation of these structures. However, a detailed understanding of the developmental progression of these structures will require further investigations. In the Supplementary Materials (Figs. S10, S11), we present hypothetical, simplified representations of the F-actin structures observed during trifurcating and bifurcating growth, organized according to their putative developmental stages. These models, however, require additional experimental validation in future studies, for instance, through experimental manipulation of F-actin (e.g., disruption via pharmacological treatments), which could help clarify the temporal relationship between actin organization and stereom formation in adult echinoderms.

Interestingly, the most commonly observed structure was the F-actin catenoid-like bridge (e.g., Fig. 6C-H and Fig. 7C), herein referred to as a “catenoid bridge”. In *Protoreaster nodosus*, catenoid bridges were observed in both galleried and diamond-type stereom, consistently forming between pairs of trabecular protrusions emerging from neighboring nodes prior to trabecular branch fusion at that site. Notably, the catenoid is a classic example of a minimal surface. We hypothesize that the tension exerted by F-actin filaments may impose a characteristic catenoid geometry, which in turn could coherently drive the formation of the stereotypical saddle-shaped morphology of trabecular surfaces. Given that saddle-shaped surfaces are a common feature across all stereom types, we further speculate that this geometric motif may be a consequence of actin-based templating governed by tension-dominated cytoskeletal dynamics. Notably, it has recently been suggested [29], although without experimental verification, that the tension of the soft tissue in sea urchins (stroma) could play a critical role in stereom morphogenesis and its saddle-shaped surfaces.

Interestingly, similar actin-mediated processes have been observed in mammalian skeletal development and in vascular morphogenesis. For example, recent work [30] has shown that the organization of F-actin in osteoblast-like cells regulates differentiation and mineralization of the extracellular matrix, highlighting the central role of the cytoskeleton in bone formation. Furthermore, it has been demonstrated [31] that VEGF (Vascular Endothelial Growth Factor) signaling involved in tubulogenesis may represent a conserved mechanism shared between spicule formation in sea urchin larvae and blood vessel development in mammals, hinting at a deep evolutionary link between biomineralization and vascular morphogenesis.

Taken together, we hypothesize that the curvature signature exhibited by the stereom of *Protoreaster nodosus* and echinoderms in general emerges from a robust growth pattern driven by the self-assembly of cytoskeletal structures. While different protein filaments may contribute to this process, F-actin appears to play a fundamental, evolutionarily conserved role, as suggested by its recurrent involvement in biomineralization across diverse lineages, including silicifying diatoms [32, 33], calcifying algae [34, 35], foraminifers [36], and echinoderms [20, this study].

## 5. Conclusions

Altogether, our findings reveal that, in the adult starfish *Protoreaster nodosus*, the formation of plates (or ossicles) bearing diamond-type stereom proceeds via two primary growth patterns, which we named trifurcation and bifurcation. These two patterns develop on distinct planes across the plate surface, although they are yet coordinated to form a coherent D-TPMS lattice. Overall, plate growth is highly dynamic, exhibiting localized variation in growth rate, which points to a high degree of cellular control over biomineralization. We demonstrate that such control is highly localized at the growing front and is likely mediated by the cytoskeleton. Indeed, irrespective of plate or stereom type, F-actin densification consistently localizes at sites of active biomineralization, particularly at elongating trabecular tips, thereby strongly supporting previous findings that underscore a critical role of the cytoskeleton in stereom morphogenesis. Furthermore, in certain regions, the formation of F-actin catenoid-like bridges precedes the development of lateral connections between adjacent stereom trabeculae, suggesting a potential role in triggering and templating trabecular fusion. We hypothesize that the cytoskeleton exerts a significant modulatory function in stereom growth, influencing also its characteristic saddle-shaped minimal surface geometry.

## Declaration of Competing Interest

The authors declare that they have no known competing financial interests or personal relationships that could have appeared to influence the work reported in this paper.

## Acknowledgements

The project was funded by NCN grant 2023/51/B/ST10/01640. Ph.D. is a Research Director of the National Fund for Scientific Research (Belgium). We thank members of the laboratory of Marine Biology at ULB for help in performing experiments. We thank the Editor and the three anonymous reviewers for their constructive and valuable comments. We also thank Paweł Bącal (Institute of Paleobiology, PAS) for his help in the acquisition of some SEM photographs.

## CRediT authorship contribution statement

Kamil Humański: Conceptualization; Formal analysis; Investigation; Methodology; Visualization; Writing - original draft; and Writing - review & editing. Giulio Facchini: Conceptualization; Formal analysis; Investigation; Methodology; Visualization; Writing - review & editing. Jenifer Croce: Conceptualization; Methodology; Writing - review & editing. Dorota Kołbuk: Formal analysis; Investigation; Methodology; Supervision; Writing - review & editing. Philippe Dubois: Methodology; Supervision; Writing - review & editing. Przemysław Gorzelak: Conceptualization; Formal analysis; Funding acquisition; Investigation; Methodology; Project administration; Supervision; Visualization; Writing - review & editing.

## Notes

### Competing Interest Statement

The authors have declared no competing interest.

### Summary of Updates

This is a revised version of the manuscript that incorporates the reviewers suggestions.

