## Supplementary material for "Morphogenesis of the diamond-type stereom microlattice and the origin of saddle-shaped minimal surfaces in the echinoderm skeleton": Supplements_Gorzelak_bioR.pdf

**Figure S1. Experimental setup with starfish *Protoreaster nodosus* during manganese labeling.**

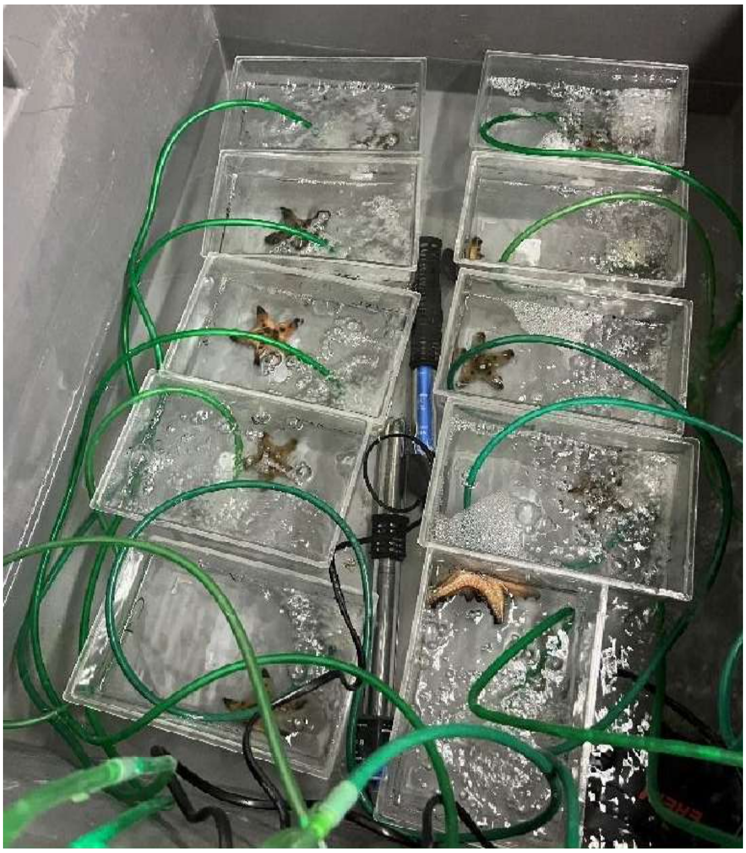

**Table S2. Seawater parameters (means and standard deviations) during the manganese labeling experiment.** Mn1-Mn5 - experimental specimens, C1-C5 - control specimens. No significant effects or interactions were detected for pH, with neither group ($p = 0.205$) nor time ($p = 0.774$) , and no significant group × time interaction was observed ($p = 0.340$). In contrast, salinity exhibited a significant effect of time within subjects ($p = 0.00549$), indicating changes across the measurement period, although no significant group effect ($p = 0.421$) or group × time interaction ($p = 0.264$) was present. Similarly, temperature showed an effect of time both between subjects ($p = 6.93 \times 10^{-5}$) and within subjects ($p = 0.00506$), but neither the main effect of group ($p = 0.540$) nor the group × time interaction ($p = 0.162$; $p = 0.933$, respectively) reached significance.

| specimen | Temperature [°C] | SD | Salinity [psu] | SD | pH | SD |
|---|---|---|---|---|---|---|
| Mn1 | 26,95 | 0,89 | 33,4 | 0,24 | 7,97 | 0,05 |
| Mn2 | 26,97 | 0,87 | 33,4 | 0,24 | 7,97 | 0,05 |
| Mn3 | 26,98 | 0,85 | 33,4 | 0,18 | 7,97 | 0,04 |
| Mn4 | 26,99 | 0,81 | 33,4 | 0,16 | 7,97 | 0,04 |
| Mn5 | 26,95 | 0,82 | 33,4 | 0,19 | 7,94 | 0,04 |
| C1 | 26,98 | 0,79 | 33,4 | 0,23 | 7,98 | 0,04 |
| C2 | 26,99 | 0,81 | 33,4 | 0,17 | 7,98 | 0,05 |
| C3 | 26,88 | 0,82 | 33,6 | 0,38 | 7,99 | 0,06 |
| C4 | 26,97 | 0,80 | 33,4 | 0,22 | 7,96 | 0,07 |
| C5 | 26,98 | 0,81 | 33,4 | 0,20 | 7,97 | 0,04 |

**Figure S3. Fragment of the arm of *Protoreaster nodosus* observed under a polarized-light microscope (crossed polars) at different rotation angles of the microscope stage**. Out of three visible plates, two larger ones, visible only partially, are composed of diamond-type stereom. The star-marked plate on the right exhibits a stereom pattern consistent with the {111} plane of the diamond-type TPMS. The 2D fast Fourier transform (FFT) in the enlarged circle (generated using ImageJ) confirms this orientation. The plate remains dark at all rotation angles, indicating that the calcite *c-axis* is perpendicular to the section plane. In contrast, the second plate, on the left, shows changes in brightness upon stage rotation, reflecting different crystallographic orientations.

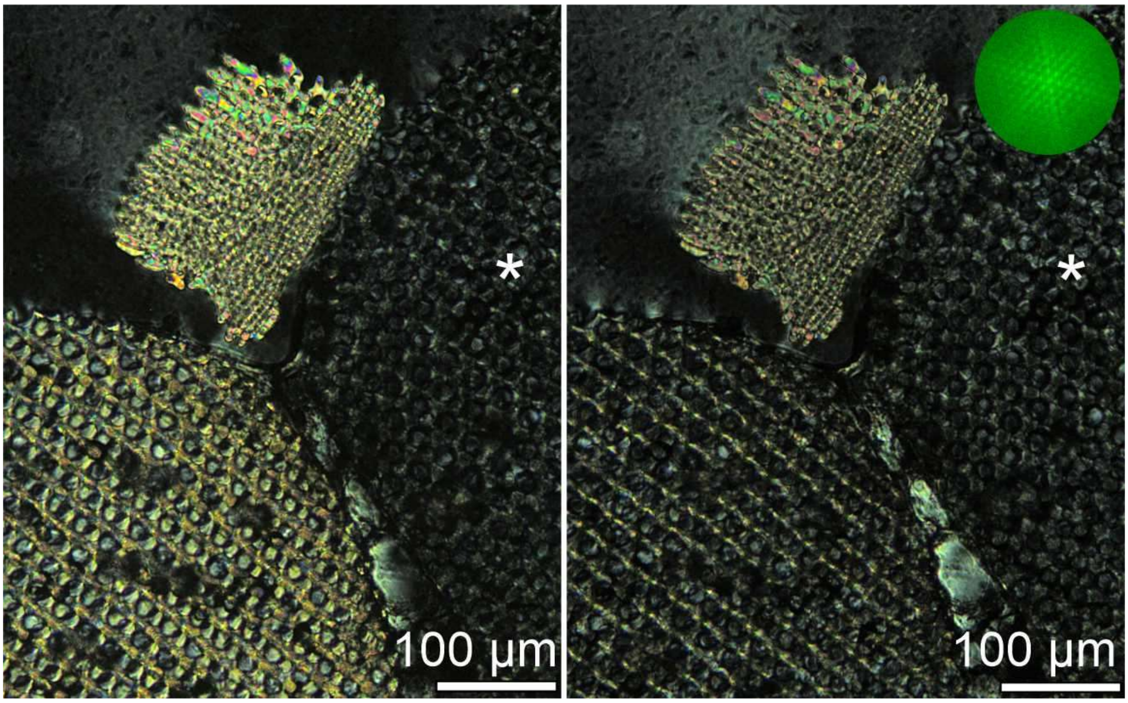


**Figure S4. Simplified cross-section of a starfish arm (A) and SEM images (B-D) showing fragments of a ventral part of a distal arm of *Protoreaster nodosus*.** In (A), different types of plates are denoted with letters: a – carinal (note that some carinal plates in *P. nodosus* grow into knob-shaped arm tubercles [Fig. S12]); b – dorsolateral; c – superomarginal; d – inferomarginal; e – adambulacral; f – ambulacral; g – spinelet. Spinelet plates are shown covering the outer surface of only a single superomarginal plate, however they are covering the outer surfaces of all plates excluding the ambulacral plates. The plates in grey are composed of labyrinthic stereom, while the plates in white are predominantly of “diamond-type” (D-TPMS) stereom, and the ones in black are of galleried stereom. In (B), SEM image showing a fragment of a ventral part of a distal arm of *Protoreaster nodosus*. Blue # and red * are adambulacral plates bearing D-TPMS stereom and spinelet plates having galleried stereom, respectively. In (C, D), zoom-ins on plates exhibiting a “diamond-type” stereom (i.e., in white plates) to illustrate the predominant node type N4 (a node connected to four branches (examples marked in red numbers 1-4)).

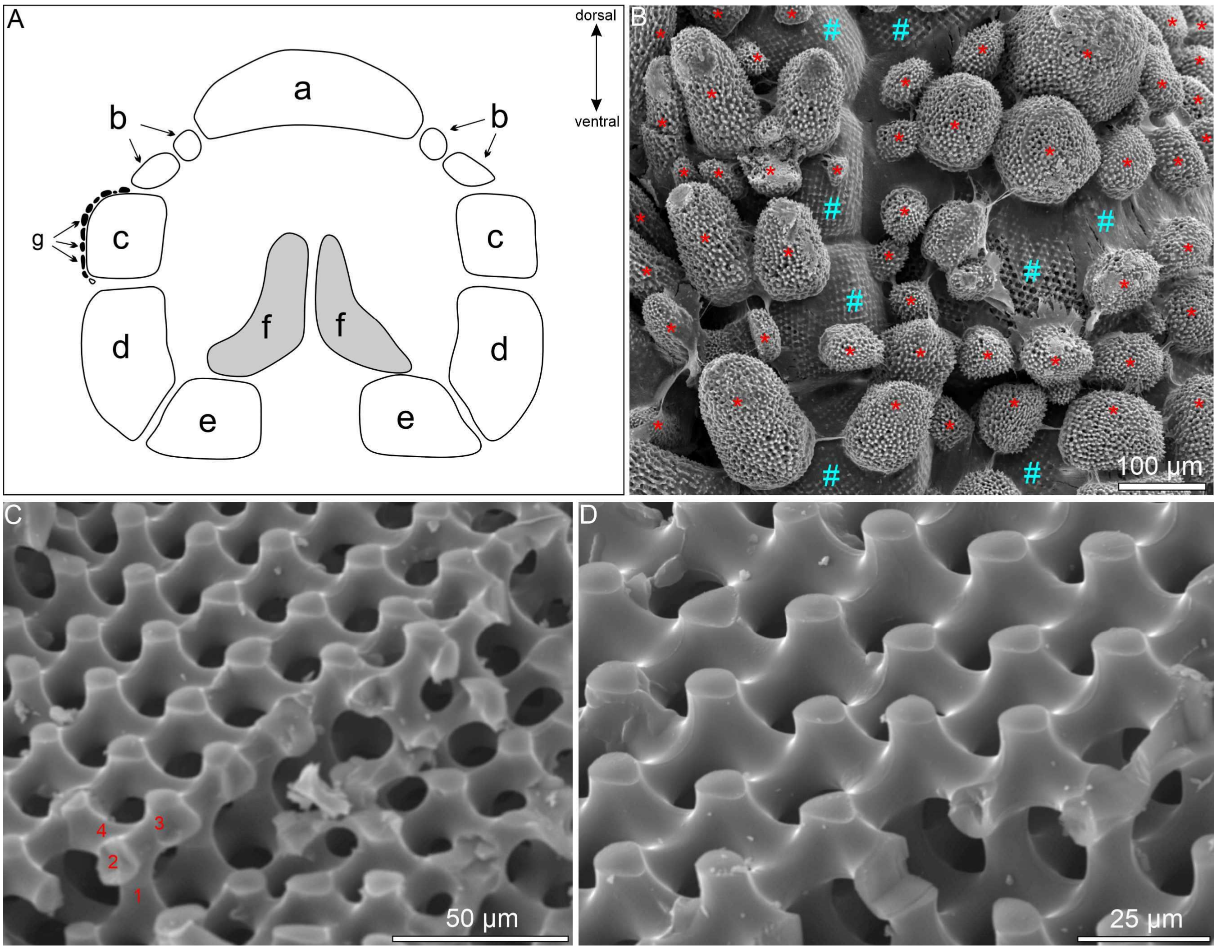


**Figure S5. Cathodoluminescence image of a carinal plate, displaying irregularities in stereom structure.** Thin section cut approximately through the sagittal plane of the plate. Manganese-induced luminescent skeleton appears as a bright orange signal. Black and dark grey regions indicate skeleton grown in normal (without $Mn^{2+}$) seawater and resin in soft tissues, respectively.

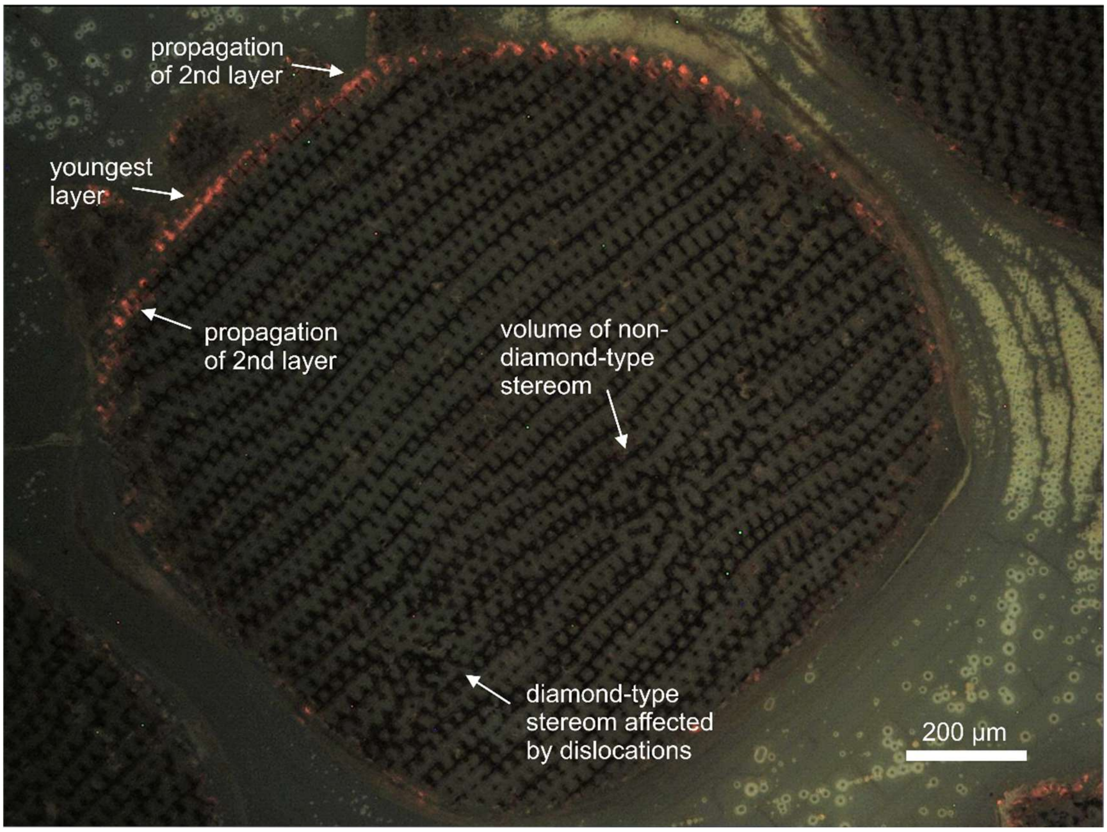


**Figure S6. Cathodoluminescence images of thin sections belonging to two specimens from the control group.** Black and dark grey regions indicate skeleton and resin in soft tissues, respectively.

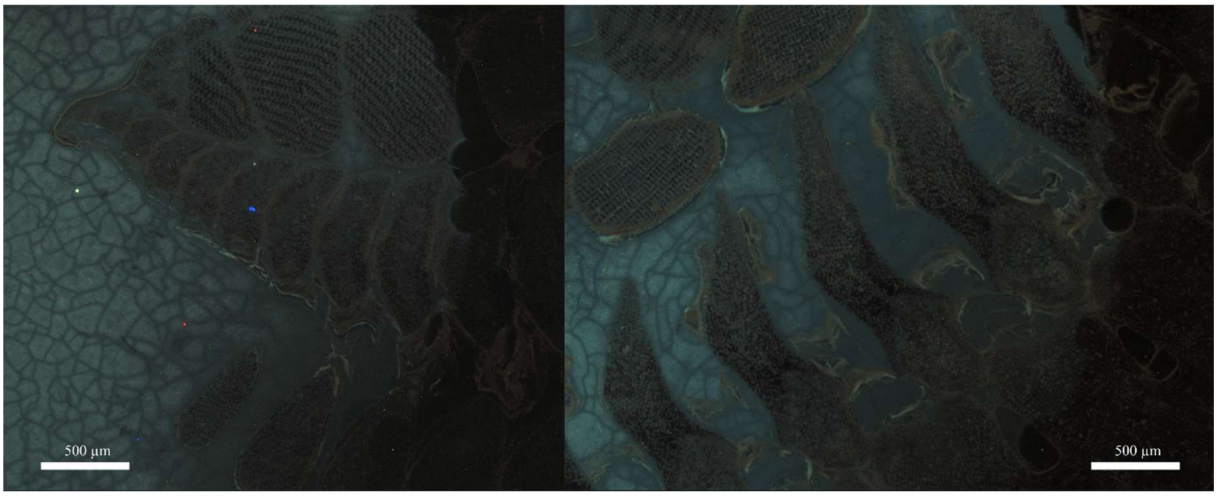

**Figure S7. Cathodoluminescence images of thin sections of individuals incubated for an extended (3-day) recovery period.** A thin white dashed line marks the outer edge of the ossicles.

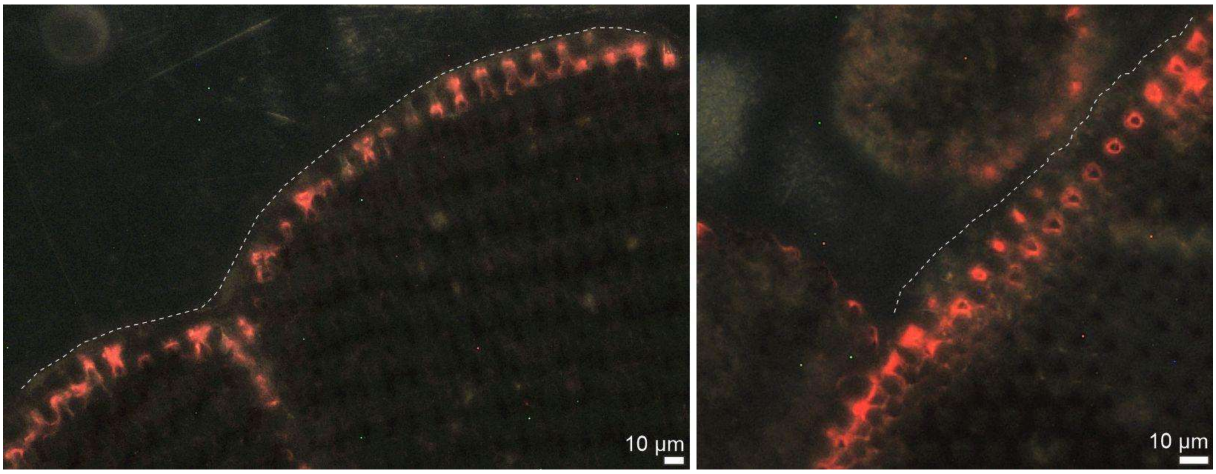


**Table S8. Rate of trifurcating layer expansion measured in five different trifurcating regions**.

| Specimen symbol, plate type | layer no. | rate of layer expansion [µm/day] |
|---|---|---|
| Mn5_1b, inferomarginal | 1 | 19,1 |
| Mn5_1b, inferomarginal | 2 | 24,4 |
| Mn5_1b, inferomarginal | 3 | 13,9 |
| Mn3_2b | 1 | 21,2 |
| Mn3_2b | 2 | 16,2 |
| Mn3_2b | 3 | 7,6 |
| Mn3_2b | 4 | 20,5 |
| Mn5_1b, inferomarginal | 1 | 15,9 |
| Mn5_1b, inferomarginal | 2 | 20,9 |

| | | |
|---|---|---|
| Mn5_1b, inferomarginal | 3 | 15,3 |
| Mn5_1b, inferomarginal | 4 | 14,8 |
| Mn3_1a, superomarginal | 1 | 12,5 |
| Mn3_1a, superomarginal | 2 | 17,9 |
| Mn3_1a, superomarginal | 3 | 13,9 |
| Mn3_1b, superomarginal | 1 | 12,2 |
| Mn3_1b, superomarginal | 2 | 17,0 |
| Mn3_1b, superomarginal | 3 | 13,1 |

**Figure S9. Cathodoluminescence (A) and SEM (B, C, D) images of superomarginal plates (A, B, C) and tubercule (D).** (A) CL image showing a section cut roughly perpendicularly to the crystallographic {111} plane of D-TPMS. (B, C) SEM images oriented along the {111} plane, either parallel to the layers (B) or slightly inclined (C) to a different crystallographic {111} plane. In any event, A, B and C images illustrate trifurcating surfaces. (D) SEM image showing bifurcation surface, where some rows are marked by thin dotted black arrows, their tips pointing in the direction of the least developed node in a row. In (A-D), nodes are shown during their growth and marked with colors and symbols corresponding to each specific stage, as in Fig. 4, for trifurcating growth (A-C), or, as in Fig. 5, for bifurcating growth (D): Red stars - Stage 1, Orange circles - Stage 2, yellow triangles - Stage 3, green rhombuses - Stage 4, blue pentagons - Stage 5, purple hexagons - Stage 6. Nodes whose stage could not be determined, as well as proto-nodes, were omitted.

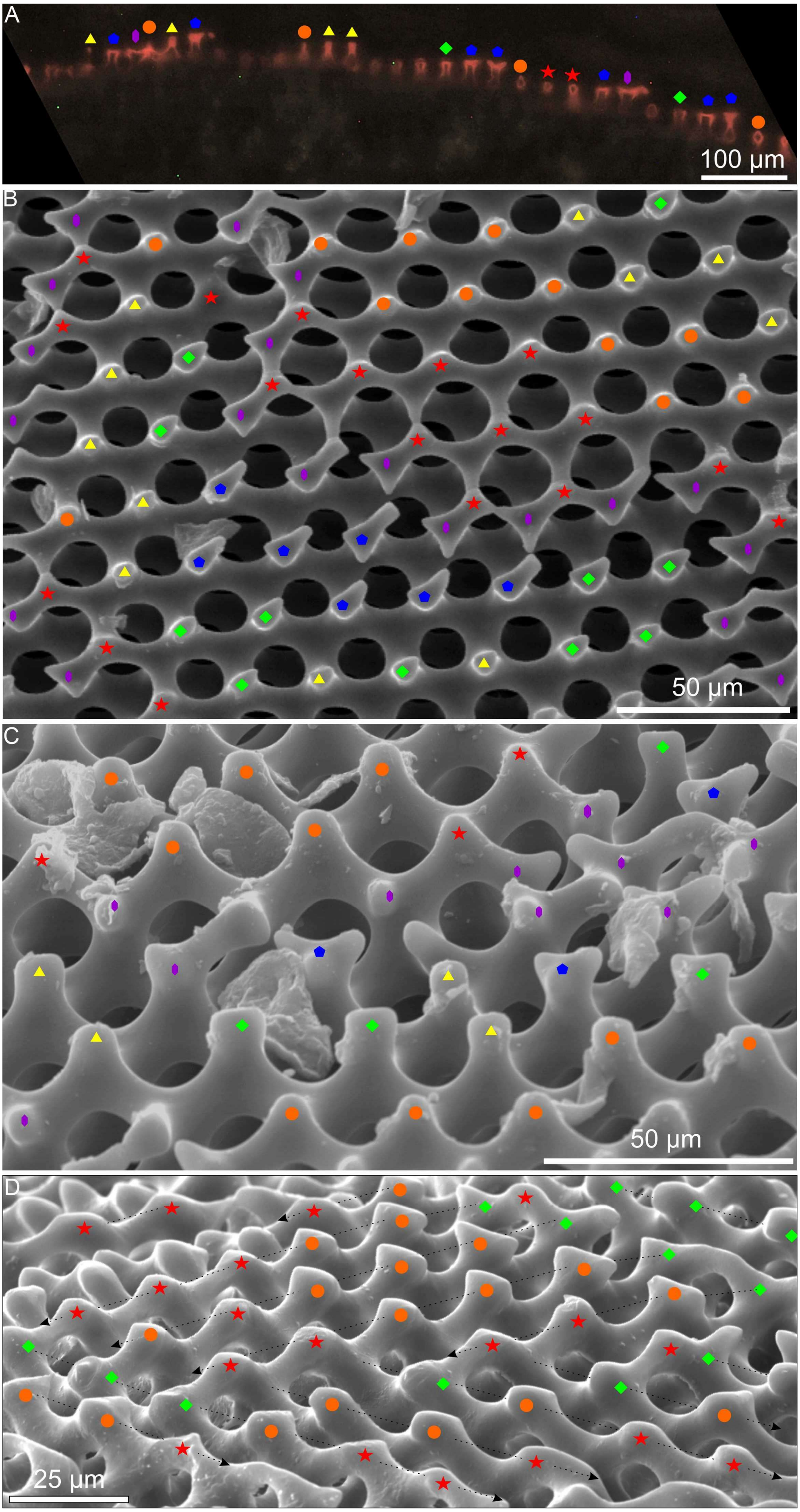
A
100 µm
B
50 µm
C
50 µm
D
25 µm

**Fig. S10. Hypothetical simplified representation of observed F-actin structures (shown on Fig. 7) associated with trifurcating growth, arranged by developmental stages of two budding nodes from a lower layer.** Node and branch thicknesses and lengths are not representative, as explained in Fig. 4. (A) Initial state, corresponding to Stage 1 in Fig. 4. (B) At the beginning of Stage 2 (as in Fig. 4) F-actin structure appears on proto-node (orientated outwards from the layer's margin), being composed of a cylindrical cap with two lateral extensions, each continuing with an F-actin filament. F-actin caps with singular filaments are observed coating budding protrusions from the layer below. (C) F-actin bridge of "pants-like" morphology forms at later Stage 2 (as in Fig. 4), connecting the proto-node with two growing branches from the lower budding nodes. Interestingly, the timing of this hypothesized connection aligns with CL and SEM observations, which indicate that around the time a crown node reaches Stage 4 of its development, protrusions budding from nearby nodes in the underlying layer begin to elongate. (D) F-actin bridge of "pants-like" morphology at Stage 3 (as in Fig. 4), connecting proto node with two new crown nodes. (E) Two catenoids after split of previous F-actin bridge at Stage 4 (as in Fig. 4). Color coding follows the scheme used in Fig. 4 plus purple – F-actin structures. The available confocal samples showed complex F-actin structures only on outward-protruding proto-nodes, while differently oriented nodes were not available for observations. This schematic is therefore not a direct replica of Fig. 4 but rather an interpretation based on the available data.

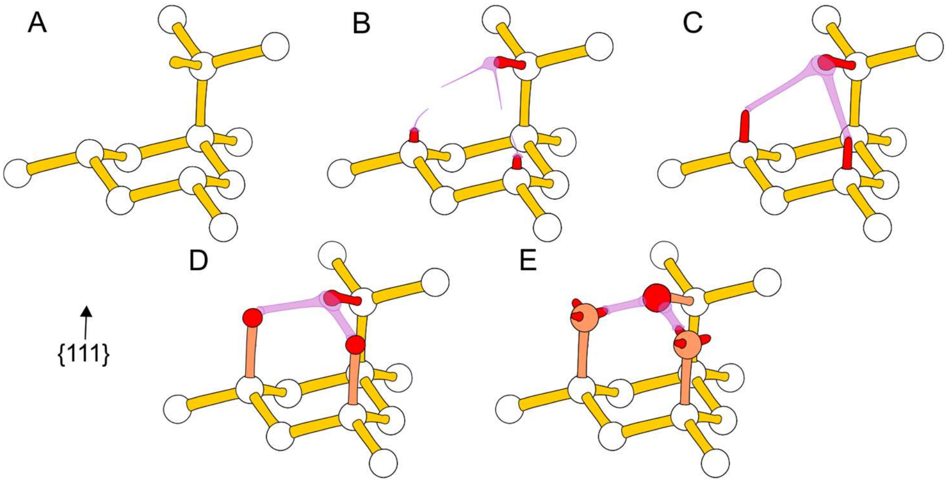


**Fig. S11. Hypothetical simplified representation of observed F-actin structures (shown on Fig. 6) associated with bifurcating growth, arranged by developmental stages of two budding nodes from a lower layer.** F-actin structures are visualized only in the region between two exemplary nodes, for which stage numbers are provided subsequently. (A) F-actin caps coating protrusions at Stage 2 (as in Fig. 5). (B) F-actin filament connecting the two cups, also at Stage 2 (as in Fig. 5). (C) A thin, elongated bridge is formed connecting the protrusion tips as they grow toward each other at Stage 3 (as in Fig. 5). (D) A minute catenoid bridge connects the two tips of the grown protrusions, slightly inclined toward one another, just prior to fusion and progressing from Stage 4 to 5 (as in Fig. 5). Purple – F-actin structures, other colors see Fig. 4.

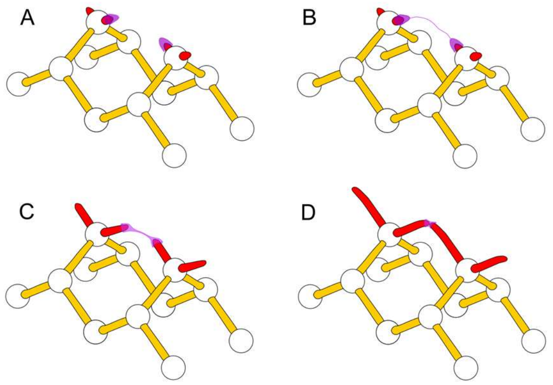

**Figure S12. SEM image of a carinal plate that grew into a knob-shaped arm tubercule, viewed from the top.** Cyan – irregular stereom filling the central top cavity, yellow – trifurcating diamond type stereom located on the rim of the cavity and followed by bifurcating stereom. Red arrows point to exemplary areas with conspicuous bifurcating growth on the sides of the tubercule, however this pattern is mixed with intermediate growth or rarely, a non-regular stereom.

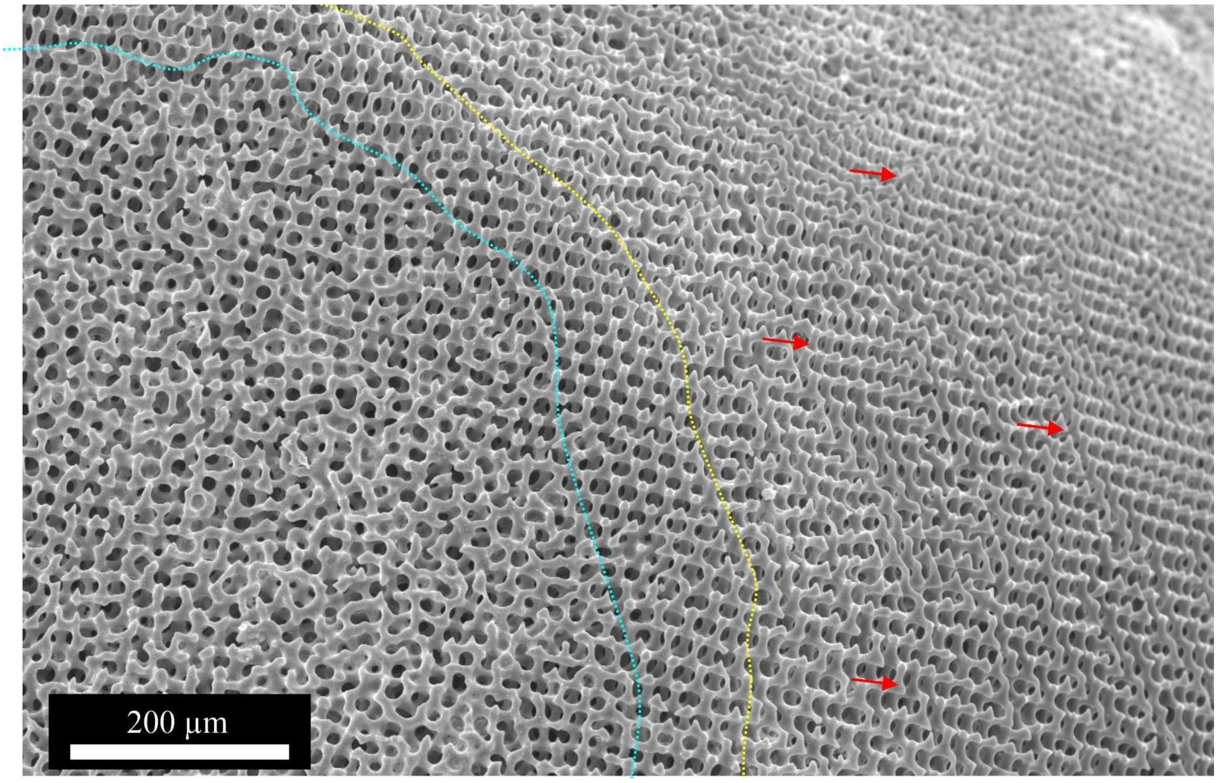
